# Western spotted skunks provide important food web linkages in forest of the Pacific Northwest

**DOI:** 10.1101/2022.04.01.486736

**Authors:** Marie I. Tosa, Damon B. Lesmeister, Jennifer M. Allen, Taal Levi

## Abstract

There are increasing concerns about the decreasing population trends of small mammalian carnivores around the world. With limited knowledge about their ecology and natural history, small mammal conservation and management remains difficult. To address one of these deficiencies for western spotted skunks (*Spilogale gracilis*), we investigated their diet in the Oregon Cascades of the Pacific Northwest during 2017 – 2019. We collected 130 spotted skunk scats opportunistically and with detection dog teams and identified prey items using DNA metabarcoding and mechanical sorting. Western spotted skunk diet consisted of invertebrates such as wasps, millipedes, and gastropods, vertebrates such as small mammals, amphibians, and birds, and plants such as Gaultheria, Rubus, and Vaccinium. Diet also consisted of items such as black-tailed deer that were likely scavenged. Comparison in diet by season revealed that spotted skunks consumed more insects during the dry season (June - August) and marginally more mammals during the wet season (September – May). We observed similar diet in areas with no record of human disturbance and areas with a history of logging. Western spotted skunks provide important food web linkages between aquatic, terrestrial, and arboreal systems by facilitating energy and nutrient transfer, and serve important functional roles of seed dispersal and scavenging. Through prey-switching, western spotted skunks may dampen the effects of irruptions of prey, such as wasps during dry springs and summers, which could then provide ecosystem resilience to environmental change.

## Introduction

Globally, many small mammalian carnivores (< 16 kg) face decreasing population trends due to multiple threats including land use change, disease, and overhunting (Belant, Schipper, and Conroy 2009; Marneweck et al. 2021). Even small carnivore species that were previously widely distributed and considered “least concern” by the IUCN, such as weasels (*Neogale* spp.), have shown signs of significant population decline (Jachowski et al. 2021). These declines are problematic because small carnivores can play important roles in ecosystem function, predator-prey dynamics, and disease transmission dynamics (Roemer, Gompper, and Van Valkenburgh 2009). Despite their potentially important roles, many small carnivores remain neglected in ecological research, data deficient, or understudied (Marneweck et al. 2021; Proulx 2010). Limited knowledge about small carnivore ecology and natural history continues to hinder management and conservation of declining populations.

In the Pacific Northwest, the western spotted skunk (*Spilogale gracilis*) is putatively a common forest carnivore, but little is known about their ecology due to their nocturnal nature (Verts, Carraway, and Kinlaw 2001) (Neiswenter and Dowler 2007; Doty and Dowler 2006; Neiswenter, Dowler, and Young 2010). Most spotted skunk literature is derived from the island spotted skunk subspecies (*S. g. amphialus*) (K. Crooks 1994b; 1994a), the congeneric eastern spotted skunk (*S. putorius*) (Lesmeister, Gompper, and Millspaugh 2009; Lesmeister et al. 2010; Kinlaw 1995) and plains spotted skunk subspecies (*S. p. interrupta*) (Crabb 1948), or other skunk species such as the pygmy skunk (*S. pygmaea*) (Cantú-Salazar et al. 2005). These spotted skunks, however, inhabit markedly different ecosystems such as on islands, prairie, or desert where there is limited forest vegetation. The Pacific Northwest, in comparison, is a temperate rainforest system that is dominated by large coniferous trees, and the functional role of western spotted skunks in this system is largely unknown. Due to their dietary plasticity, spotted skunks could vary from omnivorous generalist (e.g., eastern and plains spotted skunk) (Baker and Baker 1975; Cheeseman, Tanis, and Finck 2021; Selko 1937; Crabb 1941), insectivorous specialist (e.g., pygmy skunk) (Cantú-Salazar et al. 2005), or key carnivorous predator of small vertebrates (e.g., island spotted skunk) (K. R. Crooks and Van Vuren 1995) in Pacific Northwest forests.

Eastern and plains spotted skunk populations are in severe decline and as a result, the eastern spotted skunk is now listed as Vulnerable by the IUCN (Gompper and Jachowski 2016), and the plains spotted skunk subspecies had been petitioned for listing under the US Endangered Species Act (US Fish and Wildlife Service 2012). The mechanism for these declines are poorly understood, but multiple mechanisms have been proposed, including land-use change, disease outbreaks, or changes in predator communities (Gompper and Hackett 2005; Gompper 2017; D. Blake Sasse 2021). Although western spotted skunks are still relatively common and considered a species of Least Concern by the IUCN (Cuarón, Helgen, and Reid 2016), western spotted skunks may be prone to future, rapid declines similar to those of eastern and plains spotted skunks. Studying western spotted skunks provides an opportunity to understand the functional roles of the species and amass basic ecological knowledge that may inform conservation and land management decisions.

Land use change is a potent disturbance in the Pacific Northwest given that it is an internationally important center of timber production (Simmons et al. 2016). Forest management can influence the structure and composition of forests with unknown consequences on the ecology of western spotted skunks. One way that land use change could cause declines in spotted skunk population densities is by causing declines in prey populations. For example, potential small mammal prey such as Trowbridge’s shrews (*Sorex trowbridgii*), shrew moles (*Neurotrichus gibbsii*), red tree voles (*Arborimus longicaudus*), and flying squirrels (*Glaucomys oregonensis*), are less abundant in young forest stands than in mature and old-growth forest stands (Gilbert and Allwine 1991; Carey 1995; 1989). Disturbances such as commercial thinning can reduce small mammal prey density (Manning, Hagar, and McComb 2012). In contrast, forest management can increase the abundance of flowers and fruits of understory plants through increased light penetration onto the forest floor (Wender, Harrington, and Tappeiner 2004). Thus, it remains important to investigate how western spotted skunk diets are impacted by forest management.

Characterizing the diet of small carnivores, however, has been difficult. New techniques such as detection dogs and DNA metabarcoding have improved our ability to find scat and identify prey items, respectively. Previously, diets of spotted skunks were difficult to study because scats were often deposited in rest sites (Selko 1937; Lesmeister, Gompper, and Millspaugh 2008), not on trails, and because spotted skunks exhibit an omnivorous diet consisting of insects, small vertebrates, and fruit (Baker and Baker 1975; K. R. Crooks and Van Vuren 1995; Crabb 1941; Howell 1906). These scats were typically collected opportunistically, mechanically sorted, and morphologically identified (Ewins et al. 1994; Sándor and Ionescu 2009), but these processes had biases related to digestion that may render prey items unrecognizable (Symondson 2002; Galan, Pagès, and Cosson 2012) and misidentification of rare species (Massey et al. 2021). This can be particularly problematic for small omnivorous predators that consume a wide breadth of prey items including plants, animals, and invertebrates because identifiers must have taxonomic expertise. Moreover, misidentification of carnivores from scat morphology has been problematic and has potentially led to biased results (Morin et al. 2016). Newer genetic approaches such as DNA metabarcoding (Ji et al. 2013; Monterroso et al. 2019) can increase confidence in correctly identifying the carnivore (Morin et al. 2016), increase the number of prey items that can be correctly identified (Massey et al. 2021), and increase efficiency of identifying diet for a high volume of samples (Kartzinel et al. 2015), especially for omnivorous species (De Barba et al. 2014).

Here we use DNA metabarcoding and mechanical sorting to provide the first comprehensive analysis of western spotted skunk diet in the Pacific Northwest, and quantified seasonal variability in diet as a function of land use change. This improved understanding of western spotted skunk foraging ecology can elucidate their functional role as small vertebrate predators, insectivores, and frugivores.

## Methods

### Study Area

This study was centered around the H. J. Andrews Experimental Forest (HJA), which is located on the western slope of the Cascade Mountain Range near Blue River, Oregon (Figure 1). The is surrounded by the McKenzie River Ranger District of the Willamette National Forest. Elevations range from 410 m to 1,630 m. The maritime climate consists of warm, dry summers and mild, wet winters. Mean monthly temperatures range from 1°C in January to 18°C in July. Precipitation falls primarily as rain, is concentrated from November through March, and averages 230 cm at lower elevations and 355 cm at higher elevations (Greenland 1993; Swanson and Jones 2002). During 2018 – 2019, western Oregon experienced an extreme drought (USDM 2022). In Lane County, drought severity was greatest during August 2018 – February 2019, but abnormally dry conditions began as early as January 2018 and moderate drought conditions began as early as June 2018 (Figure S1).

**Figure 1.**
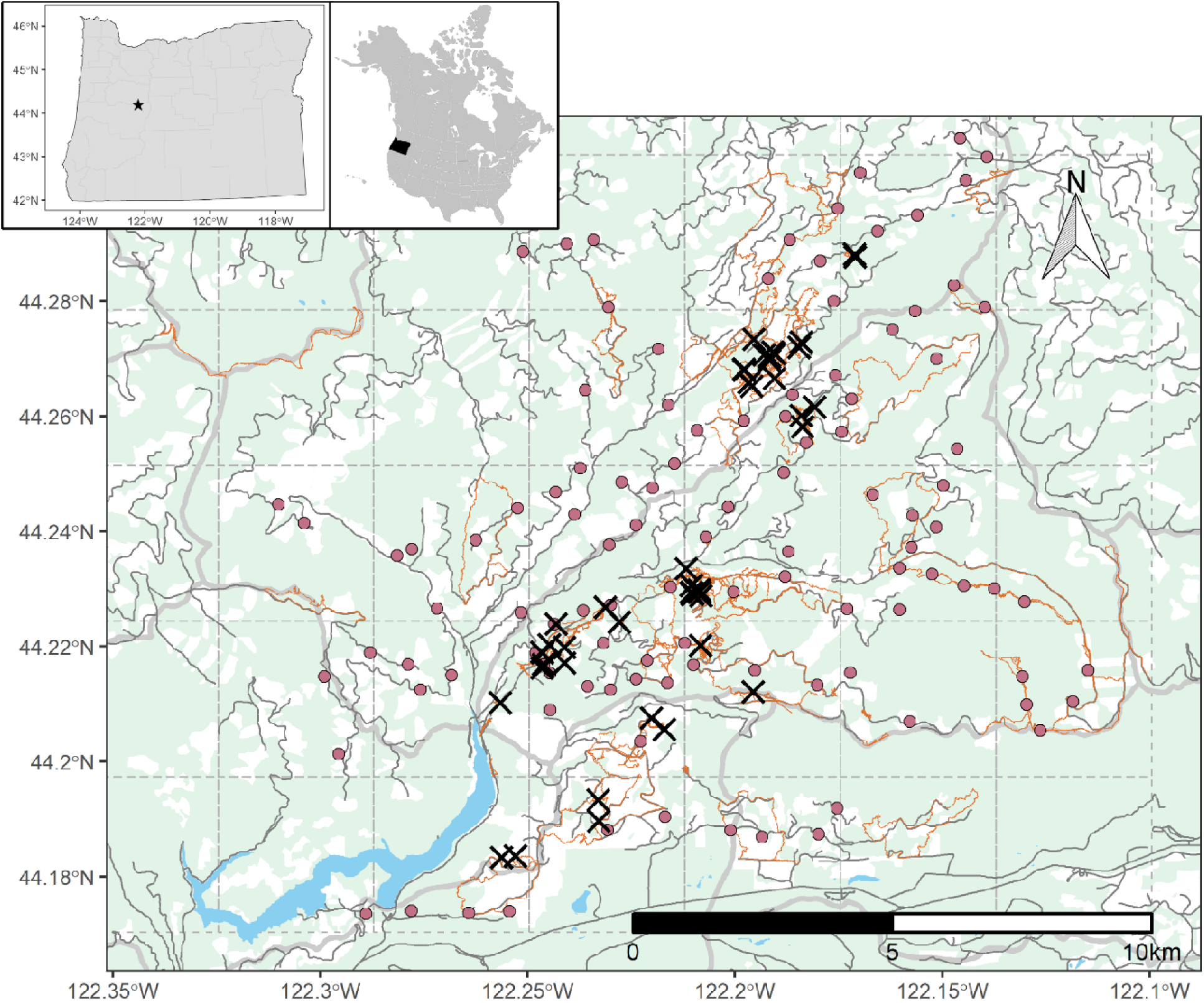
Study area within the Willamette National Forest in the Cascade Range of Oregon, USA, and locations of western spotted skunk (*Spilogale gracilis*) scats (black crosses). Detection dog tracks during the summer and fall of 2018 shown in orange, 3 x 3 km survey grids shown in grey dashed lines, and locations of camera traps shown in maroon circles. Previously logged areas shown in white. Roads shown in dark grey lines, and outlines of watersheds shown in thick light grey lines.

Lower elevation forests are dominated by Douglas-fir (*Pseudotsuga menziesii*), western hemlock (*Tsuga hetemphylla*), and western red cedar (*Thuja plicata*). Upper elevation forests are dominated by noble fir (*Abies procera*), Pacific silver fir (*Abies amabilis*), Douglas-fir, and western hemlock. The understory is variable and ranged from open to dense shrubs. Common shrubs included Oregon grape (*Mahonia aquifolium*), salal (*Gaultheria shallon*), sword fern (*Polystichum munitum*), vine maple (*Acer circinatum*), Pacific rhododendron (*Rhododendron macrophyllum*), huckleberry (*Vaccinium* spp.), and blackberry and salmonberry (*Rubus* spp.).

Before timber cutting in 1950, 65% of the HJA was covered in old-growth forest. Approximately 30% of the HJA was clear cut or shelterwood cut to create plantation forests varying in tree composition, stocking level, and age. In 1980, the HJA became a charter member of the Long Term Ecological Research network and no logging has occurred since 1985. The Willamette National Forest immediately surrounding the HJA has a similar logging history, but logging continues to occur. Currently, the HJA consists of a higher percentage of old-growth forest than the surrounding Willamette National Forest (approximately 58% in the HJA vs. 37% in the study area) (Davis et al. *In Press*). Wildfires are the primary disturbance type, followed by windthrow, landslides, root rot infections, and lateral stream channel erosion. Mean fire return interval of partial or complete stand-replacing fires for this area is 166 years and ranges from 20 years to 400 years (Teensma 1987; Morrison and Swanson 1990).

### Field methods

Our western spotted diet study was part of a larger study on their spatial ecology in the temperate rainforest ecosystem of western Oregon that was conducted between April 2017 – September 2019. During this study, we set and maintained 112 baited trail cameras and captured and tracked western spotted skunks using Tomahawk traps (Model 102 and 103, Tomahawk Live Trap Co., Hazelhurst, WI) and VHF radio-collars (M1545, 16 g; Advanced Telemetry Systems, Isanti, MN). Cameras placed in the HJA were paired with previously established long-term songbird monitoring (Frey, Hadley, and Betts 2016) and small mammal monitoring sites (Weldy et al. 2019). Cameras placed outside of the HJA were stratified based on elevation and old-growth structural index (Spies and Franklin 1988) and chosen randomly within logistical constraints. Both cameras and live traps were baited with a frozen house mouse (*Mus musculus*), a can of sardines (*Culpidae*), and/or various carnivore scent lures. We located skunks using radio-telemetry triangulation and homing techniques daily, weather permitting. Homing techniques were mainly used to locate rest site locations during the day whereas triangulation was used to locate skunks during the night when skunks were most active. All animal capture and handling were conducted in accordance with the guidelines set by the American Society of Mammalogists and were approved by the USDA Forest Service Institutional Animal Care and Use Committee (IACUC #2016-015) and the Oregon Department of Fish and Wildlife.

We collected western spotted skunk scat in multiple ways: 1) during western spotted skunk capture, 2) opportunistically while tracking western spotted skunks with radio-collars and checking trail cameras, and 3) using detection dog teams (summer and fall of 2018). Detection dog teams either surveyed 3 x 3 km grids within the study area for a minimum of 6 hours near camera trap locations where we detected western spotted skunk or focused their surveys around known spotted skunk rest sites. Focused surveys were necessary to increase scat sample sizes and increase spotted skunk scat detection rates. Moreover, western spotted skunk scats were difficult to locate opportunistically because typically, they were deposited after we tracked skunks to their rest sites, were in hard to search locations such as in hollow logs or a short distance from the rest site. We froze all scat samples until we processed them in the laboratory, and processed scats were dried for long-term storage.

### Laboratory methods

In the lab, we identified the diet of western spotted skunks using DNA metabarcoding (Massey et al. 2021; Eriksson et al. 2019) and mechanical sorting. For DNA metabarcoding, we extracted DNA in a laboratory dedicated to processing degraded DNA using the DNeasy Blood and Tissue kit (Qiagen, Germantown, Maryland) or the QIAamp Fast DNA Stool Mini Kit (Qiagen, Germantown, Maryland). We included an extraction blank with every batch of extractions as a negative control, where we used the same protocol but without a fecal sample (hereafter called extraction blanks). We kept extraction blanks throughout the DNA metabarcoding process.

Following DNA extraction, we amplified 3 regions of the mitochondria and chloroplast DNA. First, we amplified a ∼100 base-pair DNA segment of the ribosomal mitochondrial 12S gene using universal vertebrate primers (12S-V5-F’: YAGAACAGGCTCCTCTAG and 12S-V5- R: TTAGATACCCCACTATGC) (Kocher et al. 2017; Riaz et al. 2011) and the trnL gene using universal plant primers (g-F: GGGCAATCCTGAGCCAA and h-R: CCATYGAGTCTCTGCACCTATC) (Taberlet et al. 2007) in a multiplex polymerase chain reaction (PCR). In a separate singleplex PCR reaction, we amplified the COI gene using ANML universal arthropod primers (LCO1490-F: GGTCAACAAATCATAAAGATATTGG and CO1-CFMRa-R: GGWACTAATCAATTTCCAAATCC) (Jusino et al. 2019). We performed 3 PCR replicates per sample using the QIAGEN Multiplex PCR kit (Qiagen, Germantown, Maryland). To aid in identifying contamination, we performed PCR on a negative control on each plate (hereafter called PCR blanks) in addition to the extraction blanks. Each reaction was amplified with identical 8 base pair tags on the 5’ end of the forward and reverse primer that were unique to each sample to identify individual sample after pooling and to prevent misidentification of prey samples due to tag jumping (Schnell, Bohmann, and Gilbert 2015). 12S and trnL PCR reactions were carried out in a total volume of 20 μL using the following reagent mixtures: 10 μL QIAGEN Multiplex PCR Master Mix, 4 μL of forward and reverse primer mix for a final primer concentration of 200 nM, 0.2 μL of bovine serum albumin (BSA), 0.8 μL of water, and 1 μL of DNA template. After 15 minutes of initial denaturation at 95°C, we conducted 40 cycles of 94°C for 30 seconds, 58°C for 90 seconds, 72°C for 90 seconds, and a final extension at 72°C for 10 minutes. COI PCR reactions were carried out in a total volume of 20 μL using the following reagent mixtures: 4 μL of GoTaq Flexi Buffer, 1.2 μL of MgCl_2_, 0.132 μL of GoTaq Polymerase, 0.4 μL of dNTPs, 0.064 μL of BSA, 6.204 μL of water, 4 μL of each primer for a final concentration of 200 nM, and 4 μL of final DNA extract elution. After 2 minutes of initial denaturation at 95°C, we conducted 5 cycles of 94°C for 60 seconds, 45°C for 90 seconds, 72°C for 90 seconds, followed by 40 cycles of 94°C for 60 seconds, 50°C for 90 seconds, 72°C for 60 seconds, and a final extension at 72°C for 7 minutes. We normalized and pooled the PCR products and used NEBNext Ultra II Library Prep Kit (New England BioLabs, Ipswich, Massachusetts) to adapt the library pools into Illumina sequencing libraries (Illumina Inc., San Diego, California). We purified libraries using the Solid Phase Reversible Immobilization beads and sent libraries to the Center for Genome Research and Biocomputing at Oregon State University for 150 base pair paired-end sequencing on the Illumina HiSeq 3000.

We paired raw sequence reads using PEAR (Zhang et al. 2014) and demultiplexed samples based on the 8-base pair-index sequences using a custom shell script (Text S1). We counted unique reads from each sample replicate and assigned taxonomy using BLAST against the 12S, COI, and trnL sequences in a local database and GenBank (www.ncbi.nlm.nih.gov/blast). Scat amplification was considered successful if DNA sequencing produced over 100 total reads per replicate, and we limited the effects of contamination by retaining only species that consisted of more than 1% of the total reads. Furthermore, we used extraction and PCR negative controls to set additional filtering thresholds for species read counts. Species were only retained in the final species list if it was present in at least 2 of the 3 replicates and if their species distribution maps included our study area or were included on the species lists of the study area (https://andrewsforest.oregonstate.edu/about/species). We identified plants to genus since congeners are difficult to differentiate using these primers.

To mechanically sort scats, we identified remains to the lowest taxonomic order possible. We used mechanical sorting to augment results from DNA metabarcoding because of known biases introduced by mismatches in the universal invertebrate ANML primers we used, which is attributed to a lack of conserved regions across all invertebrates (Deagle et al. 2014).

We confirmed scats as defecated by western spotted skunks using the metabarcoding data following criteria: 1) western spotted skunk was the only carnivore (order: Carnivora) identified in the scat, or 2) western spotted skunk was one of the carnivores identified in the scat and the other carnivores consisted of less than 10% of the read count.

### Data analysis

We conducted analyses and produced figures using the Program R (R Core Team 2019). We quantified the importance of each taxonomic group by first calculating the frequency of occurrence of broader taxonomic group of prey (i.e., vertebrate, invertebrate, or plant). We calculated frequency of occurrence as a proportion of the number of scats in which each taxonomic group was present divided by the total number of scats. Due to the broad breadth of diet, we then calculated conditional frequency of occurrence of each species as the number of scats that contained the prey species divided by the total number of scats containing the broader taxonomic group. We also calculated the importance of each prey item by calculating the relative read abundance (RRA) for items identified through DNA metabarcoding by:

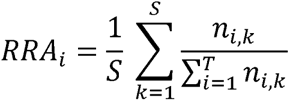

where *n_i,k_* is the number of sequences of prey species *i* in sample *k*, *S* is the number of scat samples, and *T* is the number of species. We produced figures relating taxonomy of prey items using the *metacoder* package (Foster, Sharpton, and Grünwald 2017), and we produced rarefaction curves for each taxonomic group (species for vertebrates, genus for invertebrates and plants) using the *iNEXT* package (Hsieh, Ma, and Chao 2016) to estimate completeness and expected taxonomic richness of diet based on sample size.

To investigate the effect of season and logging on western spotted skunk diet, we split data according to season (dry: June – September, wet: October – May) and location where the scat was collected (previously logged area vs. no record of logging in area). We summarized the presence or absences of each taxonomic class (e.g., Mammalia, Insecta, Gastropoda) per sample and fit a generalized linear model for multivariate data using the manyglm function in the *mvabund* package (Wang et al. 2012) to compare diet composition across season and past disturbance to the area.

## Results

During October 2017 – August 2019, we collected and genetically confirmed 130 western spotted skunk scats (n_summer_ = 47, n_fall_ = 62, n_opportunistic_ = 21). 58 scats (n_dry_ = 32, n_wet_ = 26) were collected from previously logged areas and 72 (n_dry_ = 25, n_wet_ = 47) scats were collected from areas with no record of timber harvest.

We identified 27 vertebrate species, 43 plant genera, 15 arthropod species, and 3 mollusk species as prey using DNA metabarcoding (Figure 2). The most frequent prey item (n = 15) was Atlantic herring (*Clupea harengus*), which we used to bait skunks to trail cameras and traps. Atlantic herring was the only prey item in 2 scats, so we removed these samples from the following analyses. After removing Atlantic herring, invertebrates were the most common prey items identified through metabarcoding and mechanical sorting. Invertebrates occurred in 85.2% of all scats (n = 109). Vertebrates were the next most common prey item (58.6%, n = 75), and we detected mammals in 46.9% (n = 60), birds in 14.1% (n = 18), and amphibians in 13.3% (n = 17) of all scats. Finally, we detected plants in 28.9% of all scats (n = 37).

**Figure 2.**
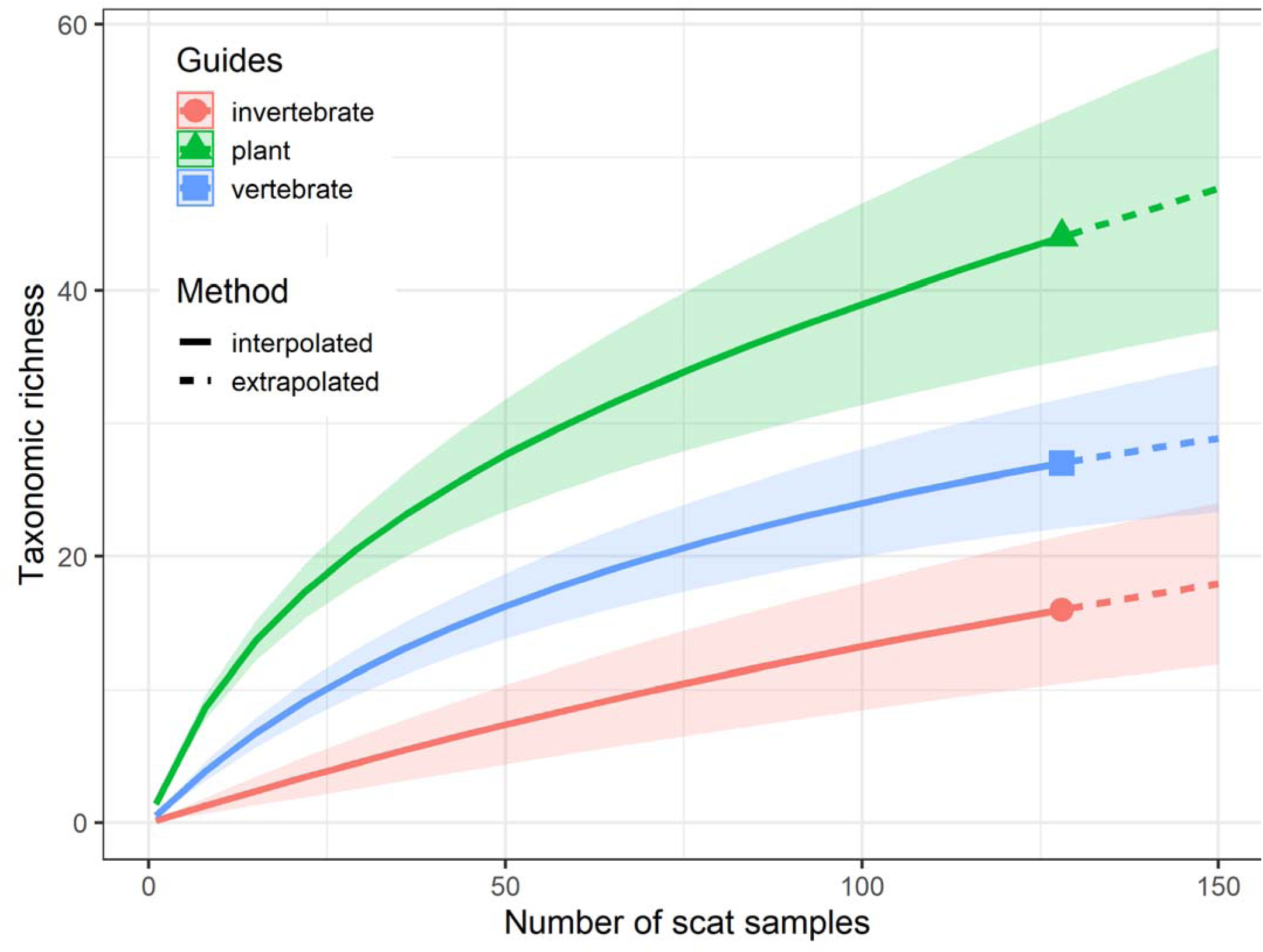
Estimation of prey taxonomic richness for western spotted skunks (*Spilogale gracilis*) in the Willamette National Forest. Vertebrate taxonomic richness (blue line) represents species richness. Invertebrate (red line) and plant (green line) richness represents genus richness.

Wasps (*Vespula* spp.) and millipedes (Diplopoda) were the top invertebrate prey items comprising 67.0% (n = 73) and 40.4% (n = 44) of scats containing invertebrates, respectively (Figure 3D). The most frequent vertebrate naturally occurring prey items were shrew mole (*Neurotrichus gibbsii*), Pacific tree frog (*Pseudacris regilla*), Townsend’s chipmunk (*Neotamias townsendii*), Swainson’s thrush (*Catharus ustulatus*), clouded salamander (*Aneides ferreus*), and Humboldt’s flying squirrel (*Glaucomys oregonensis*) (Figure 3E). The most frequent plant items were Douglas fir (*Pseudotsuga*), maple (*Acer*), Hemlock (*Tsuga*), Gaultheria (*Gaultheria*), Alder (*Alnus*), and Rhododendron (*Rhododendron*) (Figure 3F).

**Figure 3.**
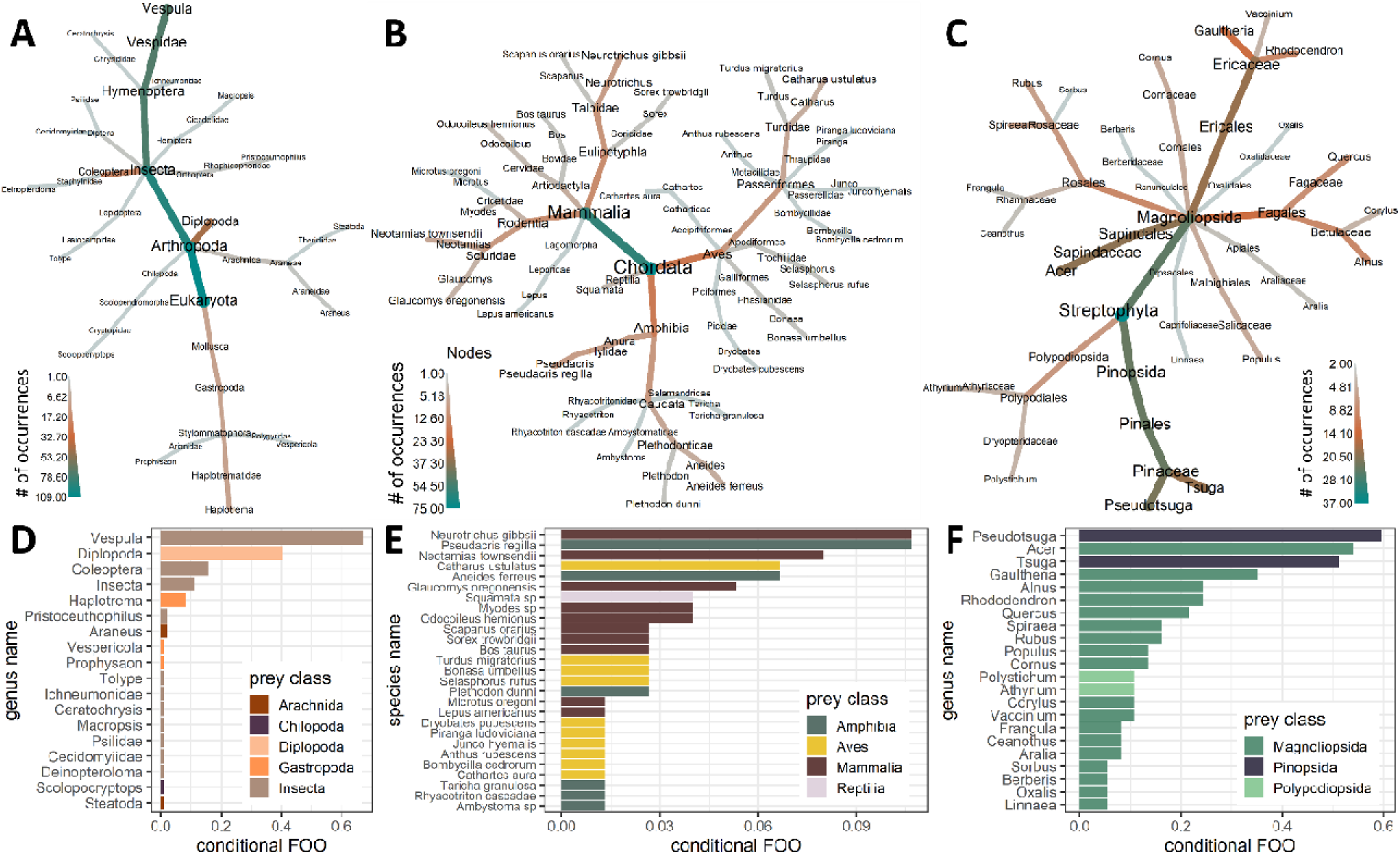
Contents of western spotted skunk (*Spilogale gracilis*) scats (n = 130) collected from 2017-2019 in the Willamette National Forest. Contents determined through DNA metabarcoding and mechanical sorting and organized by (A and D) invertebrates (n = 109), (B and E) vertebrates (n = 75), and (C and F) plants (n = 37). Taxonomic relationships of diet items shown in top row; color and size of nodes represent number of occurrences. Frequency of occurrence (FOO) conditional on presence of taxonomic group show in bottom row. Note only plants detected in more than one scat were shown.

Of the 130 scats, 51 scats (39.2%) only amplified western spotted skunk DNA and 67 scats (51.5%) did not contain any vertebrate DNA other than Atlantic herring or western spotted skunk. Manual inspection of all scat samples revealed that the majority consisted of invertebrate body parts such as wasps (Vespidae) (n = 71), millipedes (Diplopoda) (n = 43), and snail shells (Gastropoda) (n = 6) that failed to amplify with ANML primers. We also found feathers (n = 3), snake skin (n = 4), and plant material such as Douglas fir needles and bark (likely consumed incidentally along with other food items), indicating that DNA quality in some scats did not allow for detection of all diet items.

Diet composition of western spotted skunks differed based on season (LRT_season_ = 32.0, *p* = 0.001), but not based on the collection site’s logging history (LRT_logged_ = 10.7, *p* = 0.35). During the dry season, diet was composed primarily of insects, mainly wasps (LRT_insect_ = 25.4, *p* = 0.001) (Figure 4). Although plant material consumed was similar across season and the collection site’s logging history (LRT_season_ = 0.36, *p* = 0.996; LRT_logged_ = 0.007, *p* = 0.98), plants from genera that produce fruit such as Rubus (n = 6), Vaccinium (n = 1), Gaultheria (n = 10) were higher during the dry season. During the wet season, western spotted skunks consumed Gaultheria (n = 3) and Vaccinium (n = 3), but no Rubus. Similarly, although mammals and amphibians consumed were similar across season and collection site’s logging history, wet season scats consisted of more amphibians such as salamanders (*Rhyacotriton*, *Plethodon*, *Ambystoma*, and *Aneides*) and small mammals (*Neotamias townsendii*, *Neurotrichus gibbsii, Myodes* spp., *Sorex* spp.) (Figure 4).

**Figure 4.**
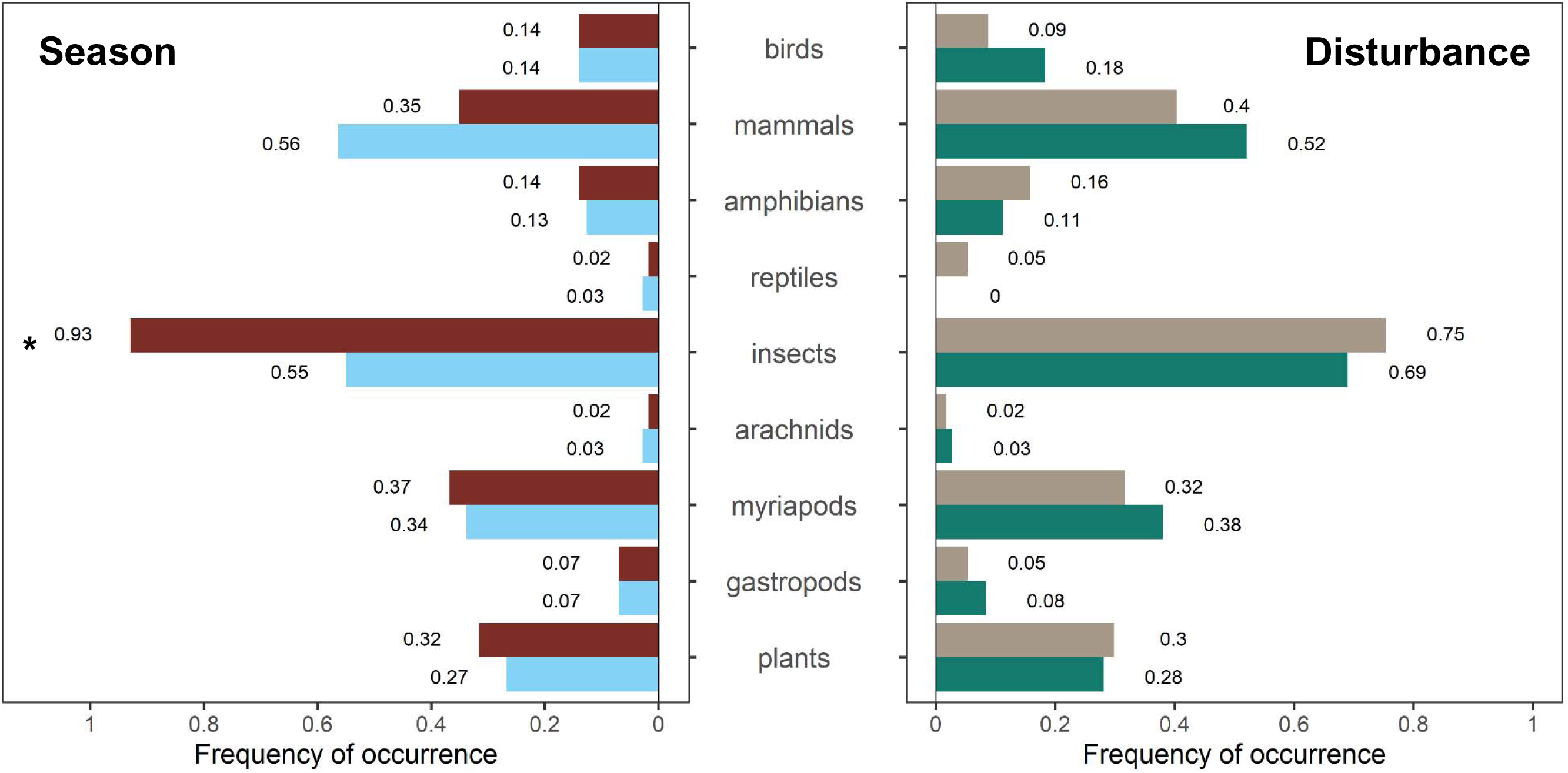
Frequency of occurrence of taxonomic groups in western spotted skunk (*Spilogale gracilis*) scats (n = 128) collected from 2017-2019 in the Willamette National Forest. Contents determined by DNA metabarcoding and mechanical sorting. Left panel shows frequency of occurrence by season (dry in red, wet in blue), right panel shows frequency of occurrence by amount of disturbance in the location we collected the scat (previously logged in tan, no record of logging in green). * represents significant differences in frequency of occurrence by taxonomic group.

## Discussion

This study provides the first data on the diet of western spotted skunks in the Pacific Northwest and represents the first use of DNA metabarcoding for high resolution spotted skunk diet analysis. In the coniferous forests of the Oregon Cascades, the western spotted skunk diet was highly diverse and included mammals, birds, amphibians, reptiles, insects, gastropods, and plants. The combined methods of DNA metabarcoding and mechanical sorting revealed that invertebrates were the primary diet items and mammals were the secondary diet item for western spotted skunks, which is consistent among other food habit studies conducted on skunks (Selko 1937; Crabb 1941; Baker and Baker 1975; Cantú-Salazar et al. 2005). The importance of these diet items shifted by season, where skunks consumed more insects during the dry season.

We detected substantially higher frequency of Vespidae than in diets reported on the island spotted skunk (K. R. Crooks and Van Vuren 1995), the pygmy skunk (Cantú-Salazar et al. 2005), and the plains spotted skunk (Howell 1906; Selko 1937; Crabb 1941). In these studies, grasshoppers and crickets (Orthoptera) and beetles (Coleoptera) were consumed more frequently (Howell 1906; Crabb 1941; Baker and Baker 1975; Cantú-Salazar et al. 2005). The scats collected for this study were collected during a year when the Oregon Cascades were unusually dry during the spring and summer (Figure S1). These conditions have been shown to be correlated with irruptions in wasp populations (Akre and Reed 1981; Dejean et al. 2011), and wasps were observed to be more abundant on the landscape (W. Gerth, personal communication). Thus, higher consumption of wasps during these irruptions suggests that western spotted skunks can switch from one prey item to another and capitalize on abundant resources as generalist predators.

As expected, the vertebrate prey base of the western spotted skunk in Oregon was much more diverse than that of the island spotted skunk that only consumed one mammal species (*Peromyscus maniculatus*) (K. R. Crooks and Van Vuren 1995) given that mammal diversity is greater in the Oregon Cascades. Unlike many of the other studies on skunks (except Howell 1906; Sprayberry and Edelman 2016), the western spotted skunk in the Oregon Cascades consumed a variety of amphibian species. DNA metabarcoding also revealed that western spotted skunks consumed 11 avian species that would have been difficult to identify with mechanical sorting. Given their generalist diet, the western spotted skunk may provide important linkages between terrestrial, aquatic, and arboreal systems in the Pacific Northwest by facilitating energy and nutrient transfer.

In addition to linking these disparate systems, western spotted skunks may perform important roles in ecosystem as seed dispersers, scavengers, and disease reservoirs. We identified plants in western spotted skunk scats with fruiting bodies including berries (e.g., Vaccinium, Rubus, Gaultheria) and mast (e.g., Acer, Quercus, Corylus). Western spotted skunks may provide key movements that allow for long-distance dispersal of seeds and may influence plant communities similar to martens and foxes (Jordano et al. 2007; González-Varo, López-Bao, and Guitián 2013). We also identified black-tailed deer in 3 scats that we collected opportunistically. We observed radio-collared western spotted skunks scavenging on kills made by mountain lions (*Puma concolor*) on trail camera videos on multiple occasions and tracked western spotted skunks to rest sites adjacent to mountain lion kill sites. This behavior has been observed in other systems (e.g., California scrub oak forest), where the western spotted skunk has the ability to displace gray fox (*Urocyon cinereoargenteus*) from carcasses and mountain lions from their kill sites (Allen, Elbroch, and Wittmer 2013; Allen et al. 2016). Furthermore, this signifies the importance of these kill sites and carrion as food resources that are worth the risks associated with being near or directly encountering larger predators (Briffa and Sneddon 2007; Allen et al. 2016; Ruprecht et al. 2021). Finally, we identified possible pathways for transmission of nematode parasites such as *Skrjabingylus* spp. which require spotted skunks as definitive hosts to complete their life cycle (Kirkland Jr. and Kirkland 1983; Higdon and Gompper 2020; LaRose, Lesmeister, and Gompper 2021; Lesmeister et al. 2008). Western spotted skunks have been shown to exhibit high prevalence and high severity of *Skrjabingylus* spp. infection (Higdon and Gompper 2020). Direct transmission of this nematode may occur through consumption of gastropods, which are the obligate intermediate host (Lankester and Anderson 1971; Kirkland Jr. and Kirkland 1983), and indirect transmission may occur through consumption of gastropod-consuming vertebrates that serve as paratenic hosts such as chipmunks *(Neotamias townsendii),* shrew moles (*Neurotrichus* gibsii), shrews (*Sorex trowbridgii*), voles *(Myodes spp.),* and amphibians (Gamble and Riewe 1982). Given that *Skrjabingylus* spp. can cause significant osteologic damage to the cranium, it is possible that this parasite could have significant impacts on individual fitness and population dynamics (Hughes et al. 2018; Lankester and Anderson 1971).

Seasonal changes in diet for western spotted skunks were similar to other skunk species, where skunks switched from consuming more insects during the dry season to more vertebrate prey during the wet season (K. R. Crooks and Van Vuren 1995; Crabb 1941). Island spotted skunks increased their consumption of crickets during the dry season and mice during the wet season (K. R. Crooks and Van Vuren 1995). Prairie spotted skunks primarily consumed rabbits and mice during the winter and spring and insects during the summer and fall (Crabb 1941). These seasonal changes in diet likely reflect changes in availability and abundance of invertebrate resources and plasticity in spotted skunk diet (Cantú-Salazar et al. 2005). In addition to the changes in availability of invertebrate resources, western spotted skunks may switch their diet to one that includes more vertebrates because they need more caloric and protein-rich input to thermoregulate and survive the harsher, colder weather (Moors 1977). Moreover, western spotted skunks breed during the fall during the wet season (Mead 1968) and, if like eastern spotted skunks (Lesmeister, Gompper, and Millspaugh 2009), males have larger home ranges when questing for mates, this could increase their energetic requirements, and therefore require an increase in caloric input.

This was the first study to examine western spotted skunk diet across scat collected from areas with different logging histories. The similarities in diet across logging history, however, is not surprising. The western spotted skunk appears to be a generalist species, and many prey species such as chipmunks may be distributed equally across these forest types. Although some of the prey species are associated with old-growth forest such as flying squirrels (*Glaucomys oregonensis*), these species still occur in logged forest, but at lower densities (Carey 1989). A relatively large portion of our study area (41.5%) is still composed of old-growth forest (Davis et al. *In Press*), and this amount of old-growth in the larger landscape may help support old-growth associated species within logged areas. Results may differ in landscapes with few remaining old-growth stands or in landscapes with more intensive logging operations. In addition, spotted skunks are highly mobile (Lesmeister, Gompper, and Millspaugh 2009) and can easily move between logged and unlogged areas in this area. Another possible explanation for similarities in diet across logged and unlogged areas is that during irruptions, species may spill over into areas that they are not typically found given lower population densities (Bock and Lepthien 1976).

Although we detected a wide variety of items in western spotted skunk scats, discrepancies between DNA metabarcoding and mechanical sorting indicated that the ANML invertebrate primers that we used poorly amplified Vespidae even though we detected Vespula in some of our samples. The potential for mismatch in the universal invertebrate primers and the biases introduced by the ANML primers we used is well-known because of a lack of conserved regions across all invertebrates (Deagle et al. 2014). Still, we used these primers because of the extensive COI reference library (Jusino et al. 2019; Elbrecht et al. 2019). This highlights the need for genera-specific primers, better universal invertebrate primers, a panel of invertebrate primers, or shotgun sequencing that does not rely on PCR to amplify target sequences so that key prey items are not missed in the future.

We detected a wide variety of plants in the western spotted skunk diet through DNA metabarcoding, but many of the genera identified may not have been consumed by western spotted skunks as a food source. When mechanically sorting scats, we discovered many intact Douglas fir needles and bark imbedded in the scat that were likely consumed incidentally, environmentally contaminated following defecation, or contaminated during scat collection. Therefore, care should be taken when interpreting some DNA metabarcoding results using plant primers.

Noticeably missing from our analysis are the fungal components of the western spotted skunk diet. Fungi are likely to contribute important nutrients (e.g., vitamins) to the western spotted skunk diet (Maser, Claridge, and Trappe 2008) and fungal dispersal by mammals is essential to plants, fungal diversity, and ecosystem function (Nuske et al. 2017). Eastern spotted skunks have been documented bringing fungal sporocarps to den sites (Sprayberry and Edelman 2016), and we have also recorded western spotted skunks carrying fungal sporocarps on trail cameras during this study. The importance of fungal diet items remains unknown for western spotted skunks and an important area of future research.

Together, understanding the diet of western spotted skunks revealed that they may be an important component of the Pacific Northwest food web. In the face of climate change, western spotted skunks may possess greater ability to withstand environmental change due to their dietary flexibility (Reed and Tosh 2019; Ducatez et al. 2020), and through their broad diet and omnivory, they may help stabilize Pacific Northwest forest food webs (Fagan 1997; Kratina et al. 2012). Through prey-switching, western spotted skunks may capitalize on, help control, and dampen the effects of irruptions of prey, such as wasps during dry springs and summers, which could then provide ecosystem resilience to environmental change. Similarly, western spotted skunks may allow species to persist and recover from low abundances through prey switching.

## Acknowledgments

We thank field technicians A. Coombs, B. Murley, K. Van Neste, and R. Rich and Rogue Detection Dogs (formerly with Conservation Canines) – H. Smith, J. Hartman, M. Poisson, C. Yee, Chester, Jack, and Scooby – for field support. We thank Levi Lab technicians for laboratory support. We also thank J. Hedges, S. Speir, and H. Thomas for mechanically sorting scats. Facilities were provided by the H. J. Andrews Experimental Forest and the Long Term Ecological Research program, administered cooperatively by the USDA Forest Service Pacific Northwest Research Station, Oregon State University, and the Willamette National Forest. Funding for this project was provided by Oregon State University, the USDA Forest Service, the National Science Foundation (LTER7 DEB-1440409), the Northwest Ecological Research Institute, and the ARCS Oregon Chapter. This publication represents the views of the authors, and any use of trade, firm, or product names is for descriptive purposes only and does not imply endorsement by the United States Government.

## Supplemental Material

### Text S1. Bioinformatics pipeline

1. Pair reads from HiSeq 3000 using PEAR

~~~
pearrun -f lane8-s001-index-ATCACG-LibA_S1_L008_R1_001.fastq -r lane8-s001-index
-ATCACG-LibA_S1_L008_R2_001.fastq -n 80 -j 12 -o ./PAIRED/SetA.pear.fastq
~~~

2. demultiplex paired reads and BLAST against NCBI database

~~~
#!/bin/bash
while read sample lib_index lib f_barcode r_barcode f_primer r_primer locus
do
SampleID=$(echo $sample"_LIB_"$lib"_FBAR_"$f_barcode"_RBAR_"$r_barcode"_LOCUS_"$locus)
         DIR=$(echo "LIB_"$lib)
         [ -d $DIR ] || mkdir $DIR
#Make reverse complements of our primer sets
         f_primer_adj=$(echo $f_primer|sed ’s/Y/[CT]/g’|sed ’s/W/[AT]/g’)
         r_primer_adj=$(echo $r_primer|sed ’s/Y/[CT]/g’|sed ’s/W/[AT]/g’)

         f_search=$f_barcode$f_primer_adj
         rcf_search=$(echo $f_barcode$f_primer_adj|rev|tr ACGT[] TGCA][)
         r_search=$r_barcode$r_primer_adj
         rcr_search=$(echo $r_barcode$r_primer_adj|rev|tr ACGT[] TGCA][)

for i in $2

         do

                   grep -oP ’(?<=’$f_search’).*(?=’$rcr_search’)’ $i| sed ’s/^/>\n/g’ >>
$DIR/$SampleID.fasta
                   grep -oP ’(?<=’$r_search’).*(?=’$rcf_search’)’ $i|tr ACGT TGCA| rev | sed
’s/^/>\n/g’ >> $DIR/$SampleID.fasta
                   cat $DIR/$SampleID.fasta | fastx_collapser > $DIR/$SampleID.clust.fasta
if [ "$locus" == "12s" ]
then

         MIDORIDB="/nfs1/FW_HMSC/Levi_Lab/Databases/MIDORIpluslocal_UNIQUE_202006 18_srRNA_SINTAX.fasta"
                   VERTMINPROB=0.8
                   usearch -threads 1 -sintax $DIR/$SampleID.clust.fasta \
                     -db ${MIDORIDB} -strand plus -sintax_cutoff ${VERTMINPROB} \
                            -tabbedout $DIR/$SampleID.clust.fasta.usearch

                   blastn -db /nfs1/FW_HMSC/Levi_Lab/Databases/Marten.Nov2019.Blast.fasta \
                   -query $DIR/$SampleID.clust.fasta \
                   -outfmt "6 qseqid sseqid sscinames staxids pident qcovs evalue bitscore qseq
sseq" \
                   -max_target_seqs 1 -evalue 1e-5 \
                   >> $DIR/$SampleID.clust.fasta.assigned

                   blastn -db /nfs1/FW_HMSC/Levi_Lab/Databases/nt_12s_eukaryotes \
                   -query $DIR/$SampleID.clust.fasta \
                   -outfmt "6 qseqid sseqid sscinames staxids pident qcovs evalue bitscore qseq
sseq" \
                   -max_target_seqs 1 -evalue 1e-5 \
                   >> $DIR/$SampleID.clust.fasta.assigned

                   awk -F "\t" ’$8 > maxvals[$1] {lines[$1]=$0 ; maxvals[$1]=$8}END { for (tag in lines) print lines[tag] }’ $DIR/$SampleID.clust.fasta.assigned | sort -nk1 >
$DIR/$SampleID.clust.fasta.assigned.bestmatch
elif [ "$locus" == "COI" ]
then
MIDORIDBCOI="/nfs1/FW_HMSC/Levi_Lab/Databases/MIDORI_UNIQUE_20180221_COI_SINTA X.fasta"
        INVERTMINPROB=0.8
        usearch -threads 1 -sintax $DIR/$SampleID.clust.fasta \
            -db ${MIDORIDBCOI} -strand plus -sintax_cutoff ${INVERTMINPROB} \
              -tabbedout $DIR/$SampleID.clust.fasta.usearch

                   blastn -db /nfs1/FW_HMSC/Levi_Lab/Databases/nt_COI_eukaryotes \
                   -query $DIR/$SampleID.clust.fasta \
                   -outfmt "6 qseqid sseqid sscinames staxids pident qcovs evalue bitscore qseq
sseq" \
                   -max_target_seqs 1 -evalue 1e-5 \
                   >> $DIR/$SampleID.clust.fasta.assigned

                   cat $DIR/$SampleID.clust.fasta.assigned | sort -nk1 >
$DIR/$SampleID.clust.fasta.assigned.bestmatch

elif [ "$locus" == "ITS" ]
then
ITS="/nfs1/FW_HMSC/Levi_Lab/Databases/sh_general_release_dynamic_s_02.02.2019.fasta"
           ITSMINPROB=0.8
           usearch -threads 1 -sintax $DIR/$SampleID.clust.fasta \
             -db ${ITS} -strand plus -sintax_cutoff ${ITSMINPROB} \
             -tabbedout $DIR/$SampleID.clust.fasta.usearch
                   blastn -db
/nfs1/FW_HMSC/Levi_Lab/Databases/sh_general_release_dynamic_s_02.02.2019.fasta \
             -query $DIR/$SampleID.clust.fasta \
             -outfmt "6 qseqid sseqid sscinames staxids pident qcovs evalue bitscore qseq sseq" \
             -max_target_seqs 1 -evalue 1e-5 \
             >> $DIR/$SampleID.clust.fasta.assigned

          cat $DIR/$SampleID.clust.fasta.assigned | sort -nk1 >
$DIR/$SampleID.clust.fasta.assigned.bestmatch

elif [ "$locus" == "trnL" ]
then
                  blastn -db /nfs1/FW_HMSC/Levi_Lab/Databases/trnL_Kwhite.fasta -task
blastn-short \
                  -query $DIR/$SampleID.clust.fasta \
                  -outfmt "6 qseqid sseqid sscinames staxids pident qcovs evalue bitscore qseq
sseq" \
                  -max_target_seqs 1 -evalue 1e-5 \
                  >> $DIR/$SampleID.clust.fasta.ALASKA.assigned
          cat $DIR/$SampleID.clust.fasta.ALASKA.assigned | sort -nk1 >
$DIR/$SampleID.clust.fasta.ALASKA.assigned.bestmatch

                  blastn -db /nfs1/FW_HMSC/Levi_Lab/Databases/trnLsequences_nospace.fasta -
task blastn-short \
                  -query $DIR/$SampleID.clust.fasta \
                  -outfmt "6 qseqid sseqid sscinames staxids pident qcovs evalue bitscore qseq
sseq" \
                  -max_target_seqs 1 -evalue 1e-5 \
                  >> $DIR/$SampleID.clust.fasta.trnLnew.assigned
        cat $DIR/$SampleID.clust.fasta.trnLnew.assigned | sort -nk1 >
$DIR/$SampleID.clust.fasta.trnLnew.assigned.bestmatch

else

                  blastn -db nt \
                  -query $DIR/$SampleID.clust.fasta \
                  -task blastn -outfmt "6 qseqid sseqid sscinames staxids pident qcovs evalue
bitscore qseq sseq" \
                  -max_target_seqs 1 -evalue 1e-5 \
                  >> $DIR/$SampleID.clust.fasta.assigned

        cat $DIR/$SampleID.clust.fasta.assigned | sort -nk1 >
$DIR/$SampleID.clust.fasta.assigned.bestmatch

fi

                  done

done<$1
~~~

**Figure S1.**
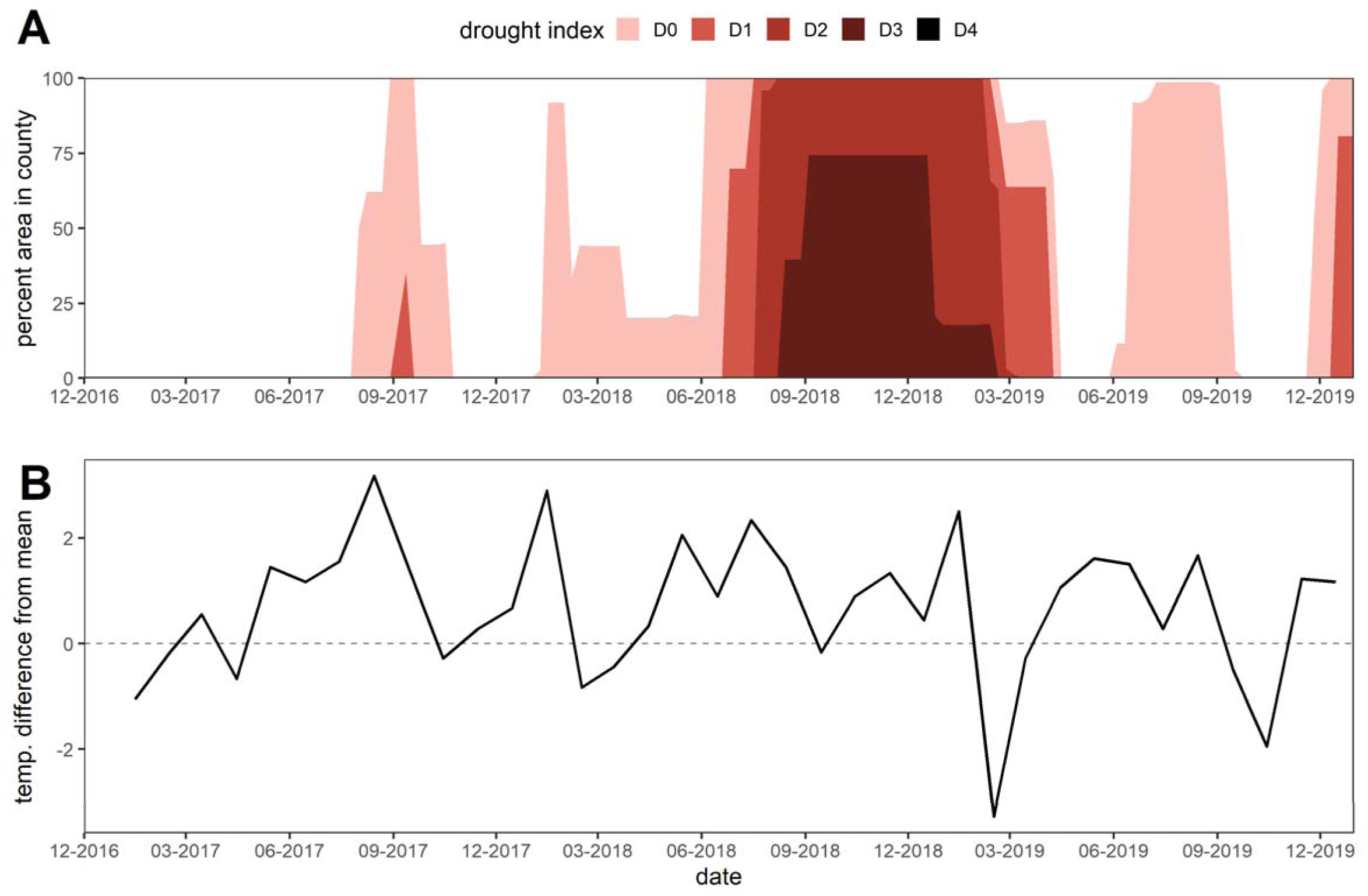
Weekly climate values for Lane County, Oregon during 2017 – 2019. (A) Values indicating the percentage of county in drought categories of abnormally dry (D0), moderate drought (D1), severe drought (D2), extreme drought (D3) and exceptional drought (D4). (B) Mean temperature difference (°C) compared to mean monthly temperature calculated from data from 1901 – 2000.

**Figure S2.**
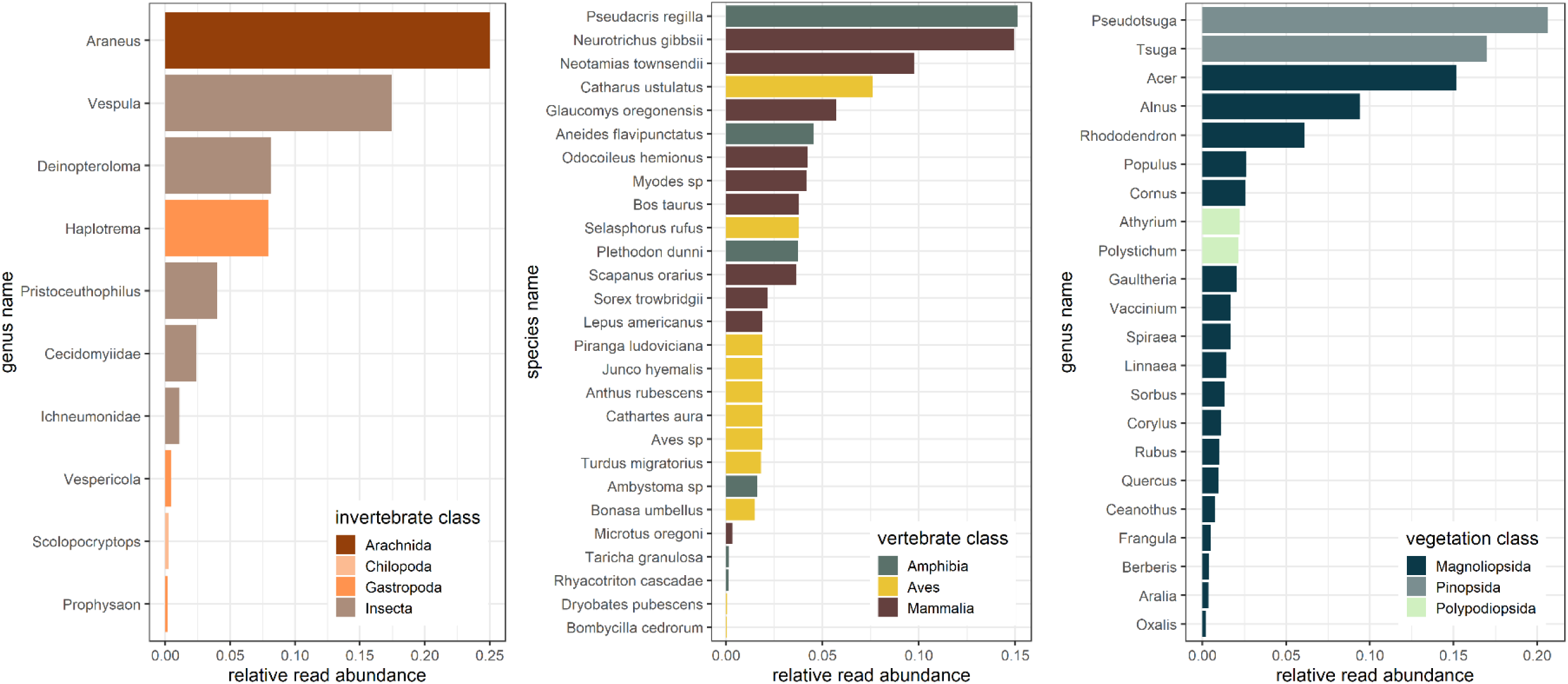
Relative read abundances of (A) invertebrates, (B) vertebrates, and (C) plants in western spotted skunks (*Spilogale gracilis*) diets during 2017-2019 in the Willamette National Forest near Blue River, Oregon. Note figure only represents diet items identified through DNA metabarcoding.

**Figure S3.**
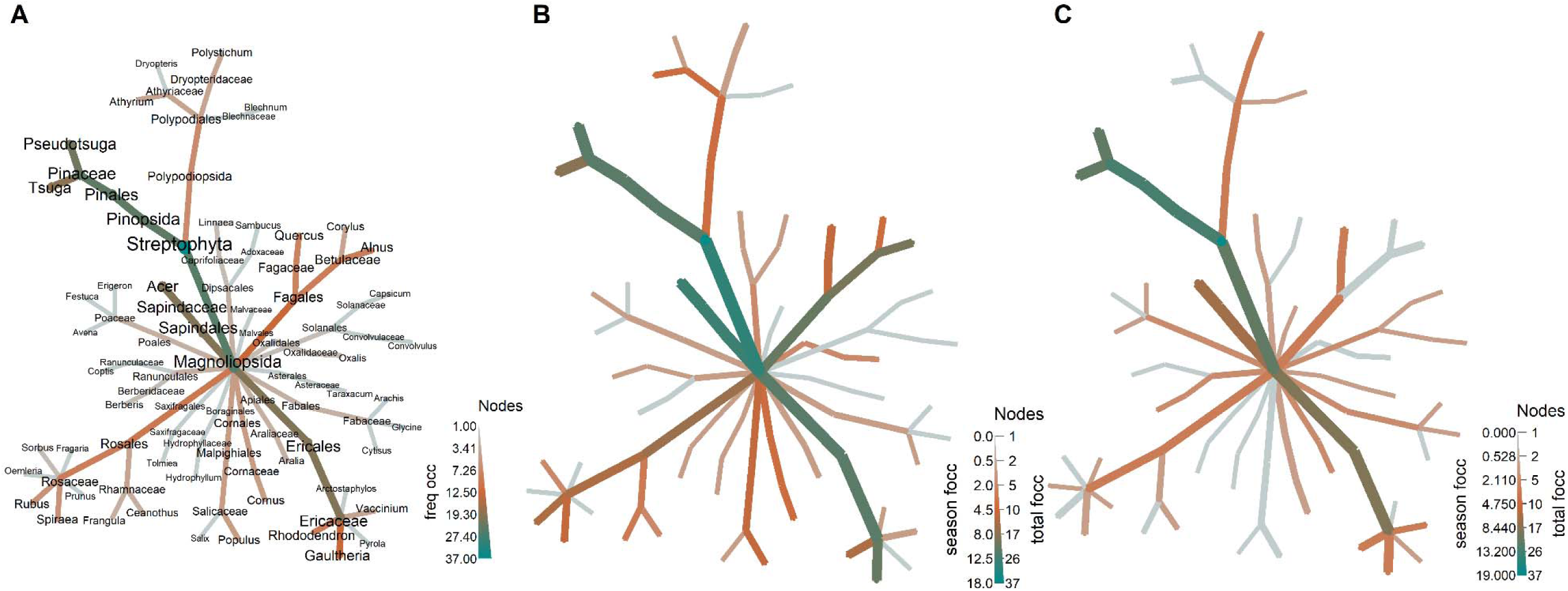
Plant diet of western spotted skunks (*Spilogale gracilis*) identified through DNA metabarcoding. (A) Plant identified in all scats collected from 2017-2019 (n = 37), (B) plants identified in scats collected during the dry season (n = 18), and (C) plants identified in scats collected during the wet season (n = 19) in the Willamette National Forest.

## Notes

### Competing Interest Statement

The authors have declared no competing interest.

### Summary of Updates

Species ID of Aneides has been corrected to a species that occurs within the study area. Aneides flavipunctatus to Aneides ferreus.

