## Supplemental Material for "Western spotted skunks provide important food web linkages in forest of the Pacific Northwest"

This material includes:

**Text** S1. Bioinformatics pipeline

Figure S1**Figure S1.** Weekly climate values for Lane County, Oregon during 2017 – 2019. (A) Values indicating the percentage of county in drought categories of abnormally dry (D0), moderate drought (D1), severe drought (D2), extreme drought (D3) and exceptional drought (D4).

cat $DIR/$SampleID.clust.fasta.assigned | sort -nk1 > $DIR/$SampleID.clust.fasta.assigned.bestmatch

fi

done

done<$1

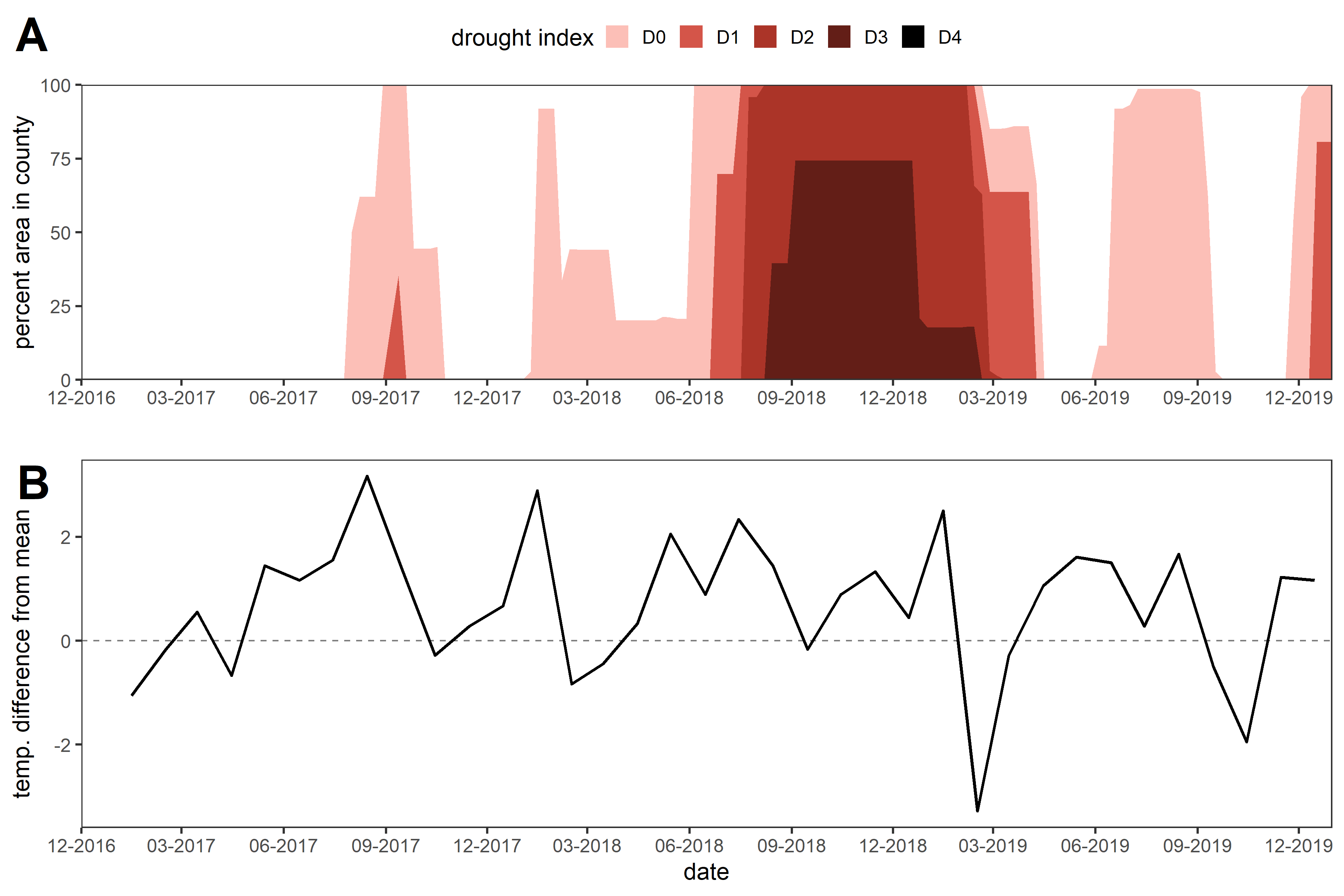

Figure S1. Weekly climate values for Lane County, Oregon during 2017 – 2019. (A) Values indicating the percentage of county in drought categories of abnormally dry (D0), moderate drought (D1), severe drought (D2), extreme drought (D3) and exceptional drought (D4). (B) Mean temperature difference (°C) compared to mean monthly temperature calculated from data from 1901 – 2000.

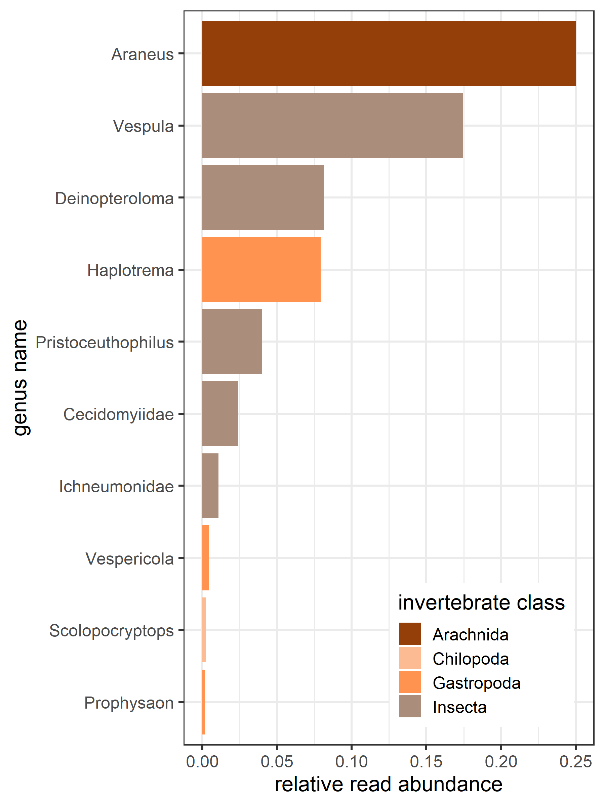

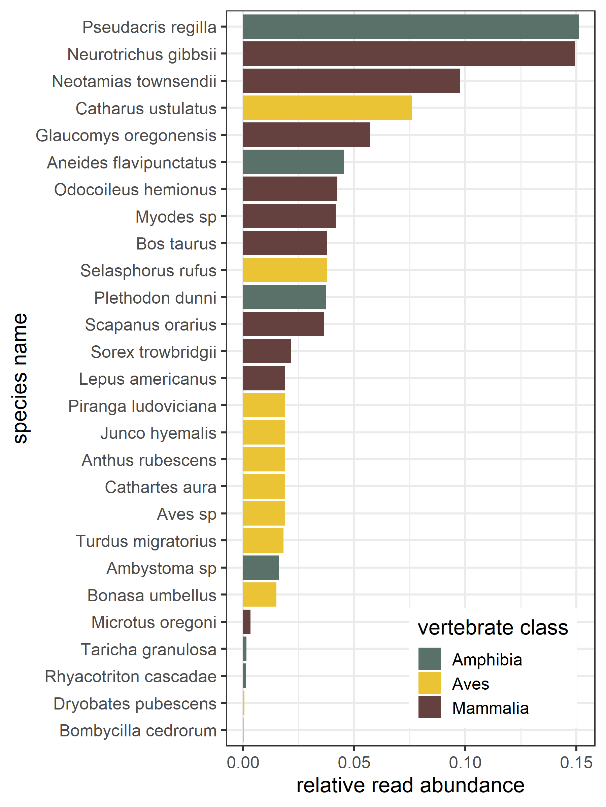

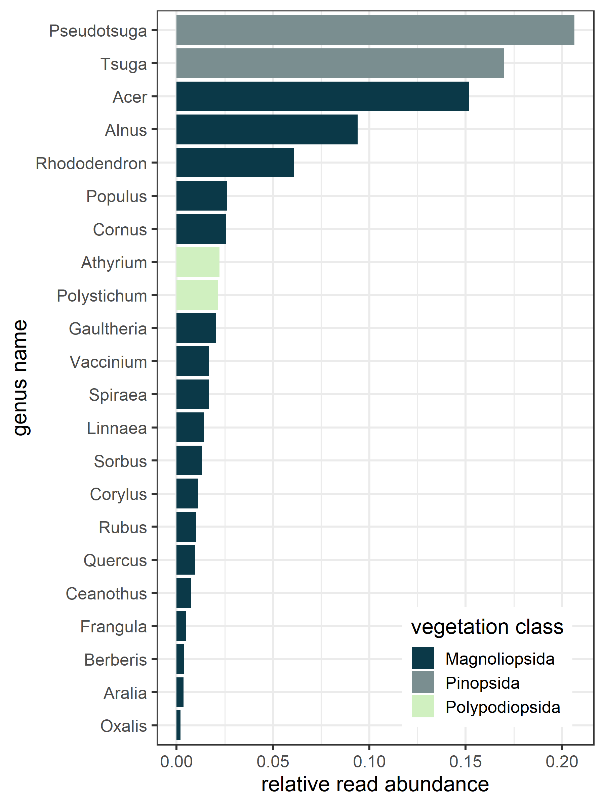

Figure S2. Relative read abundances of (A) invertebrates, (B) vertebrates, and (C) plants in western spotted skunks (*Spilogale gracilis*) diets during 2017-2019 in the Willamette National Forest near Blue River, Oregon. Note figure only represents diet items identified through DNA metabarcoding.

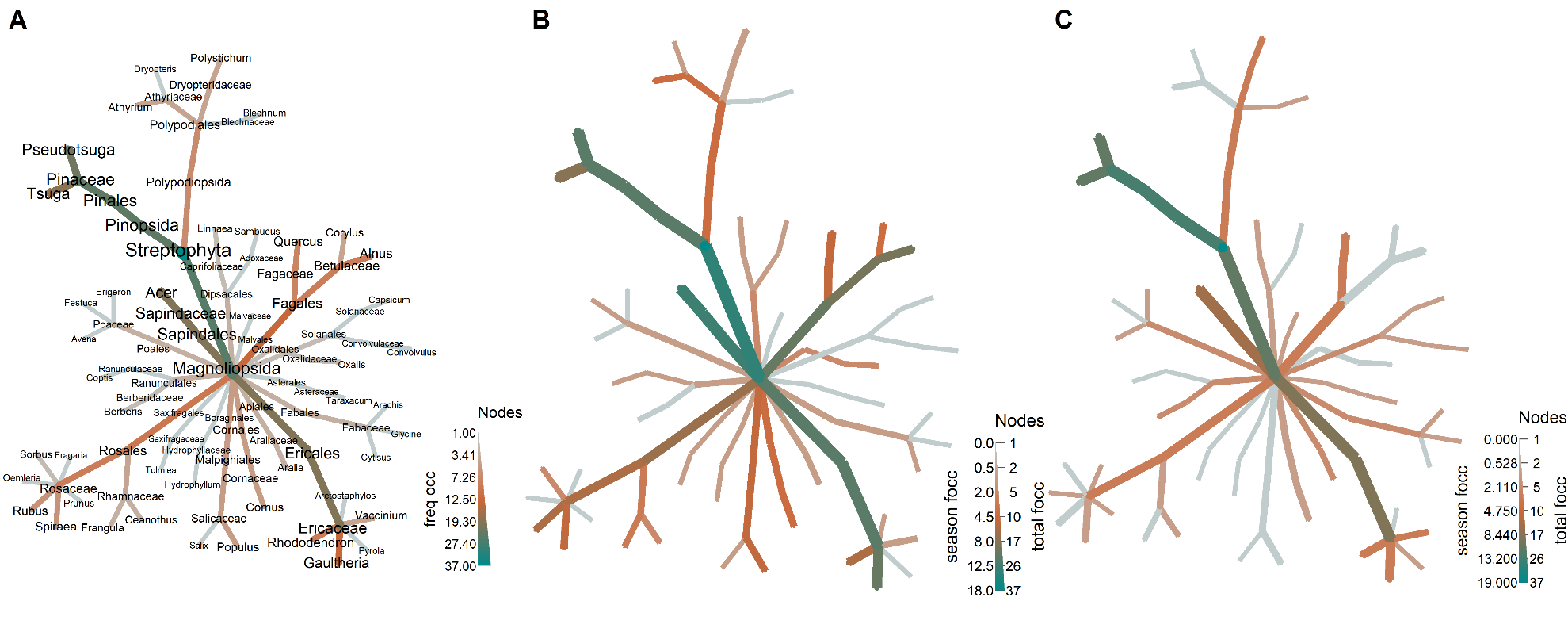

Figure S3. Plant diet of western spotted skunks (*Spilogale gracilis*) identified through DNA metabarcoding. (A) Plant identified in all scats collected from 2017-2019 (n = 37), (B) plants identified in scats collected during the dry season (n = 18), and (C) plants identified in scats collected during the wet season (n = 19) in the Willamette National Forest.
